# The cell cycle gene regulatory DREAM complex is disrupted by high expression of oncogenic B-Myb

**DOI:** 10.1101/199539

**Authors:** Audra N. Iness, Jessica Felthousen, Varsha Ananthapadmanabhan, Keelan Z. Guiley, Mikhail Dozmorov, Seth M. Rubin, Larisa Litovchick

## Abstract

The oncogene *MYBL2* (encoding B-Myb) is a poor prognostic biomarker in many cancers. B-Myb interacts with the MuvB core of five proteins (LIN9, LIN37, LIN52, LIN53/RBBP4, and LIN54) to form the MMB (Myb-MuvB) complex and promotes expression of late cell cycle genes necessary for progression through mitosis. Both *MYBL2* amplification and over-expression are associated with deregulation of the cell cycle and increased cell proliferation. Alternatively, by interacting with E2F4-DP1 and p130 or p107, the MuvB core becomes part of the DREAM complex (DP, RB-like, E2F, and MuvB). The DREAM complex opposes MMB by globally repressing cell cycle genes in G0/G1, maintaining the cell in a quiescent state. However, the specific mechanism by which B-Myb alters the cell cycle is not well understood. Our analysis of The Cancer Genome Atlas data revealed significant upregulation of DREAM and MMB target genes in breast and ovarian cancer with *MYBL2* gain. Given that most of the DREAM target genes are not directly regulated by B-Myb, we investigated the effects of B-Myb on DREAM formation. We found that depletion of B-Myb results in increased DREAM formation in human cancer cells, while its overexpression inhibits DREAM formation in the non-transformed cells. Since the MuvB core subunit LIN52 is essential for assembly of both the DREAM and MMB complexes, we tested whether B-Myb disrupts DREAM by sequestering LIN52. Overexpression of LIN52 did not increase either DREAM or MMB formation, but instead increased the turnover rate of the endogenous LIN52 protein. Interestingly, co-expression of B-Myb increased the expression of both endogenous and overexpressed LIN52 while knockdown of B-Myb had an opposite effect. We found that regulation of LIN52 occurs at the protein level, and that activity of DYRK1A kinase, the enzyme that triggers DREAM complex formation by phosphorylating LIN52, is required for this regulation. These findings are the first to implicate B-Myb in the disassembly of the DREAM complex and offer insight into the underlying mechanisms of poor prognostic value of *MYBL2* amplification in cancer. We conclude that B-Myb mediates its oncogenic effects not only by increasing mitotic gene expression by the MMB complex, but also by broad disruption of cell cycle gene regulatory programs through compromised DREAM formation.

## Introduction

Since its discovery in 1988, several studies have established *MYBL2* (encoding B-Myb) as a clinically important oncogene. ^1, 2^ Indeed, *MYBL2* is part of the commercially available Onco*type* DX^®^ screening panel and validated DCIS (Ductal Carcinoma *in situ*) Score^™^, used to clinically predict the risk of local recurrence in patients with breast cancer. ^3, 4^ Although B-Myb is a well-recognized biomarker, particularly in breast cancer, the specific cellular mechanisms of its oncogenic activities are not fully understood.

Previous studies characterized B-Myb as a transcription factor involved in cell cycle regulation, present in almost all proliferating cells. ^5^ Furthermore, B-Myb is essential for cell proliferation, as evidenced by failure of inner cell mass formation and early embryonic death of *MYBL2* knock-out mice. ^6^ Deregulation of cell cycle gene expression is one of B-Myb’s oncogenic properties. ^7-9^ Studies in *Drosophila* and human cells revealed that B-Myb regulates transcription as part of evolutionarily conserved multi-subunit protein complexes sharing common subunits with DNA-binding complexes formed by retinoblastoma (RB) family members. ^10^ In *Drosophila*, Myb regulates expression of developmental and cell cycle genes by associating with the dREAM (RB, E2F, and Myb) complex. This complex also includes five proteins homologous to the products of the *C. elegans* multi-vulva class B (MuvB) genes: LIN9, LIN37, LIN52, LIN53/RBBP4 and LIN54.^11, 12^ In the mammalian cell cycle, the DREAM (DP, RB-like, E2F, and MuvB) complex is important for modulating gene expression by repressing more than 800 cell cycle genes (including *MYBL2*) in quiescent cells. ^13^ It is assembled in G0/G1 when the MuvB subunit, LIN52, is phosphorylated at serine-28 (S28) by DYRK1A (dual-specificity tyrosine-phosphorylation regulated kinase 1A), bringing together RB-like p130 and the MuvB core of five proteins. ^14^ Cell cycle progression occurs when p130 is phosphorylated by cyclin-dependent kinases and dissociates from MuvB. ^15^ MuvB then binds B-Myb in the S phase, forming the MMB (<u>M</u>yb-<u>M</u>uv<u>B</u>) complex. MMB formation is necessary for G2/M gene expression by FOXM1 and is therefore associated with increased mitotic activity. ^16, 17^ Importantly, LIN52 is required for the MuvB core to bind B-Myb, making LIN52 an essential component of both the DREAM and MMB complexes. ^15^ Unlike the S28 phosphorylation requirement for DREAM assembly, LIN52 binding to B-Myb is phosphorylation-independent. Notably, a recent study found that MMB complex formation is necessary for lung tumor formation and drives cellular proliferation through expression of mitotic kinesins. ^8^ Additionally, several MMB downstream target genes are included in a chromosomal instability signature (CIN) used as a predictor of outcomes in multiple cancer types. ^18-20^ Despite these findings, the specific mechanism of deregulated cell cycle gene expression by B-Myb remains elusive.

Studies of DREAM disruption show that DREAM-deficient cells are unable to maintain quiescence and have MMB complex levels comparable to proliferative cells, despite signals for cell cycle arrest. ^21^ The phenotype of DREAM disruption thus closely resembles that of B-Myb over-expression. ^22, 23^ These results support the idea of mandatory MMB complex formation when DREAM is unable to assemble and also suggest that B-Myb, when over-expressed, plays a causal role in disrupting DREAM. When DREAM is inactivated by genetic mutations in mice, abnormal binding of B-Myb to MuvB during the G1 phase of the cell cycle correlates with loss of DREAM target gene repression, leading to cell cycle deregulation. ^21^ Since *MYBL2* is a target gene suppressed by DREAM, it is difficult to determine whether or not B-Myb over-expression in cancer disrupts DREAM or if DREAM disruption occurs by other mechanisms, then leading to enhanced B-Myb expression. Therefore, we investigated the regulation of DREAM by B-Myb as a potential mechanism for the cell cycle defects observed in cancers with high B-Myb levels. We show that increased expression of B-Myb disrupts DREAM by recruitment of LIN52 to the MMB complex, compromising repression of DREAM-targeted cell cycle genes. These findings implicate global cell cycle deregulation as a means by which B-Myb exerts its oncogenic effects.

## Results

### B-Myb regulates DREAM

#### High B-Myb level is associated with compromised DREAM formation and function

To test our hypothesis that B-Myb disrupts DREAM, we first looked at microarray data from The Cancer Genome Atlas (TCGA) to determine the relationship between B-Myb level and cell cycle gene expression. We chose to focus on high grade serous ovarian carcinoma (HGSOC), since it exhibits a high prevalence of *MYBL2* amplification or gain (55%) and represents a cancer type in which B-Myb is associated with a poor prognosis (**Fig. 1A, 1S**). ^24^ Similar analyses were conducted in parallel for breast cancer, for which the significance of B-Myb overexpression is well-documented (**Fig. 1S**). ^25-31^ As expected, we found that high B-Myb levels are associated with increased MMB target gene expression. Furthermore, expression of DREAM target genes were also significantly upregulated in the tumor samples with high B-Myb expression (**Fig. 1C**), and the top 50 most differentially expressed genes in this cohort have been previously annotated as DREAM target genes (**Fig. 1D, 1S**). ^13^ Given these results, we used SKOV3 serous ovarian carcinoma cells (with *MYBL2* gene amplification) to investigate mechanisms of how B-Myb modulates DREAM complex formation. ^32^ In asynchronously growing SKOV3 cells, DREAM exists at low steady state levels and RNAi-mediated knock-down of B-Myb enhances DREAM formation (**Fig. 2A**). RT-qPCR analysis revealed decreased expression of representative DREAM target genes (*FOXM1* and *CCNB2*) upon B-Myb knockdown, in agreement with the observed increase of the DREAM formation (**Fig. 2B**). ^33-35^

**Figure 1:**
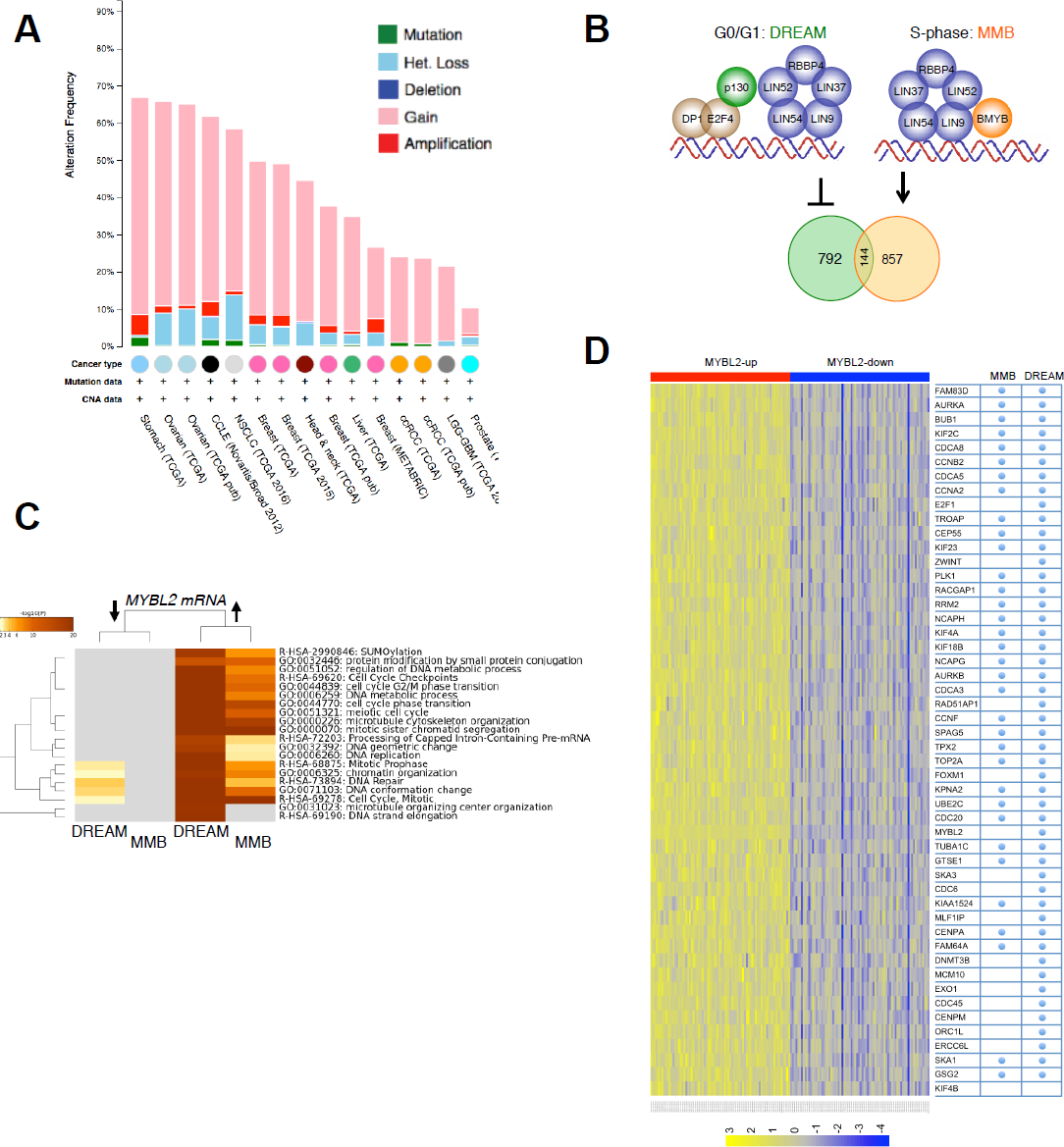
DREAM and MMB target genes are upregulated in the presence of high B-Myb. (A) Analysis of copy number variations of *MYBL2* gene across different cancer types reveals widespread gains. Graph shows studies with more that 300 cases visualized using cBio.org. **(B)** Schema of the DREAM (repressor) and MMB (activator) complexes that share a common MuvB core (blue circles). Venn diagram shows the numerical breakdown of known DREAM-and MMB-specific as well as shared target genes identified by global location analysis. **(C)** DREAM and MMB target genes are significantly upregulated in ovarian cancers with high B-Myb expression (Fisher’s exact test p-value = 0.0065). **(D)** Annotation of the top 50 up-regulated genes in TCGA ovarian cancer samples with high B-Myb expression shows significant enrichment of the DREAM and MMB target genes (X^2^ with Yates correction < 0.001).

**Figure 2:**
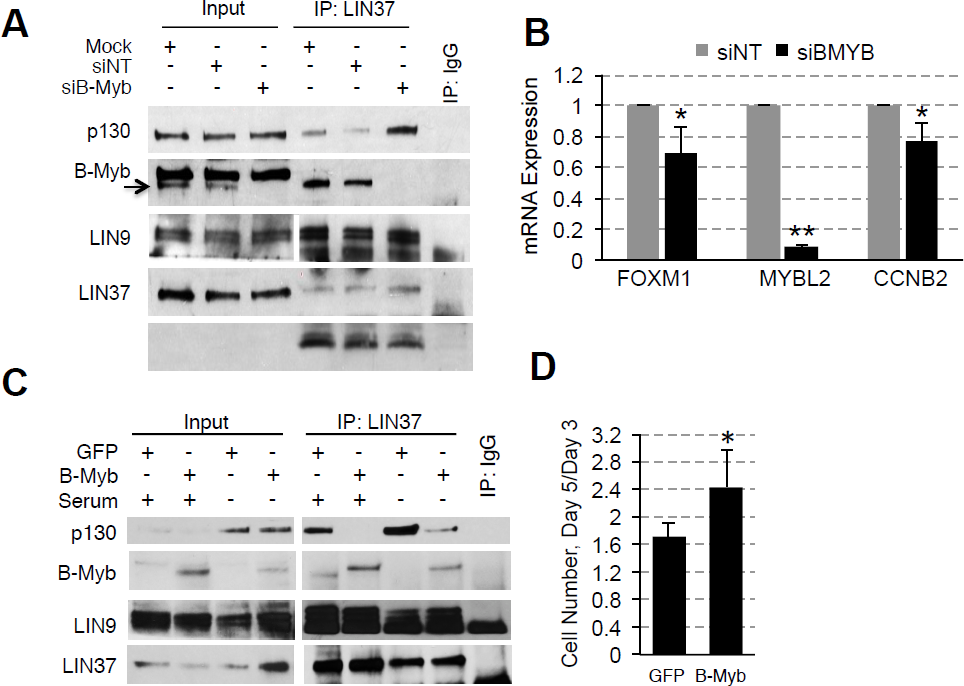
B-Myb expression levels influence DREAM formation and function. (A) IP/WB analysis shows increased binding of p130 to LIN37 (indicative of DREAM formation) in SKOV3 cells transfected with B-Myb-specific siRNA compared to control cells. **(B)** RT-qPCR analysis shows decreased expression of *FOXM1* (DREAM target) and *CCNB2* (DREAM and MMB target) genes upon B-Myb knock-down in SKOV3 cells. raph shows average ± stdev of three independent experiments (asterisk, Student’s t-test p<0.05). **(C)** IP/WB analysis of the DREAM complex assembly in BJ-hTERT fibroblasts stably expressing HA-GFP (control) or HA-B-Myb. Where indicated, cells were incubated without serum for 48 h to induce G0/G1 arrest and DREAM complex formation. **(D)** Increased proliferation of BJ-hTERT cell line expressing HA-B-Myb compared to HA-GFP (control). Graph shows average increase of cell number on day 5 relative to day 3 after plating measured in three independent experiments using ATPlite assay. Asterisk, Student’s t-test p<0.05.

The effects of B-Myb gain of function on DREAM formation were subsequently assessed by expressing HA-B-Myb in non-transformed human foreskin fibroblasts immortalized with hTERT (BJ-hTERT). We hypothesized that high B-Myb levels could impede DREAM assembly, thus overexpression of B-Myb should suppress DREAM regardless of environmental signals favoring DREAM formation. Indeed, in the cells that were serum-starved to induce cell cycle arrest and DREAM assembly, less p130 co-immunoprecipitated with a shared MuvB complex protein, LIN37, in the presence of B-Myb overexpression as compared to cells expressing the control vector (HA-GFP) (**Fig. 2C**). Consistent with this finding, we also observed a significantly greater proliferation rate of HA-B-Myb cells compared to control cells (**Fig. 2D**). Collectively, the data support a model in which B-Myb, when present at high levels, interferes with DREAM formation and function, altering cell cycle gene expression programs. Furthermore, evidence of MMB formation under conditions that normally favor DREAM assembly suggests that B-Myb achieves DREAM disruption by binding to MuvB.

### LIN52 regulation

#### LIN52 is regulated at the protein level

Since LIN52 is required for the interaction between either B-Myb or p130 with MuvB core, we next wanted to determine if increasing availability of LIN52 by ectopic overexpression could rescue DREAM formation in the presence of HA-B-Myb. Therefore, we transduced BJ-hTERT fibroblasts expressing HA-B-Myb, (shown in **Figure 2C**), with retrovirus to express LIN52-V5. Control BJ-hTERT cells as well as cells expressing HA-B-Myb alone, or together with LIN52-V5, were treated with the CDK4/6 inhibitor, palbociclib, to encourage DREAM formation. ^15^ As shown in **Figure 3A,** DREAM assembly was enhanced with palbociclib treatment in control cells, but not in the cell lines with ectopic expression of HA-B-Myb. At the same time, inappropriate MMB formation was also observed in the presence of palbociclib treatment. Interestingly, these effects persisted in BJ-hTERT cells in which LIN52-V5 was co-expressed with HA-B-Myb even with palbociclib treatment, suggesting that highly expressed B-Myb could sequester LIN52 (and MuvB) away from p130 under these conditions. It is also possible that the overall availability of the MuvB core did not increase upon ectopic expression of LIN52. In support of this conclusion, the amount of co-precipitated LIN9 remained the same as in control cells despite more LIN52 co-immunoprecipitated with LIN37 from LIN52-V5-expressing cells. Furthermore, the amount of the endogenous LIN52 bound to LIN37 in the presence of LIN52-V5 was markedly reduced, indicating that availability of other subunits could limit the formation of MuvB.

**Figure 3:**
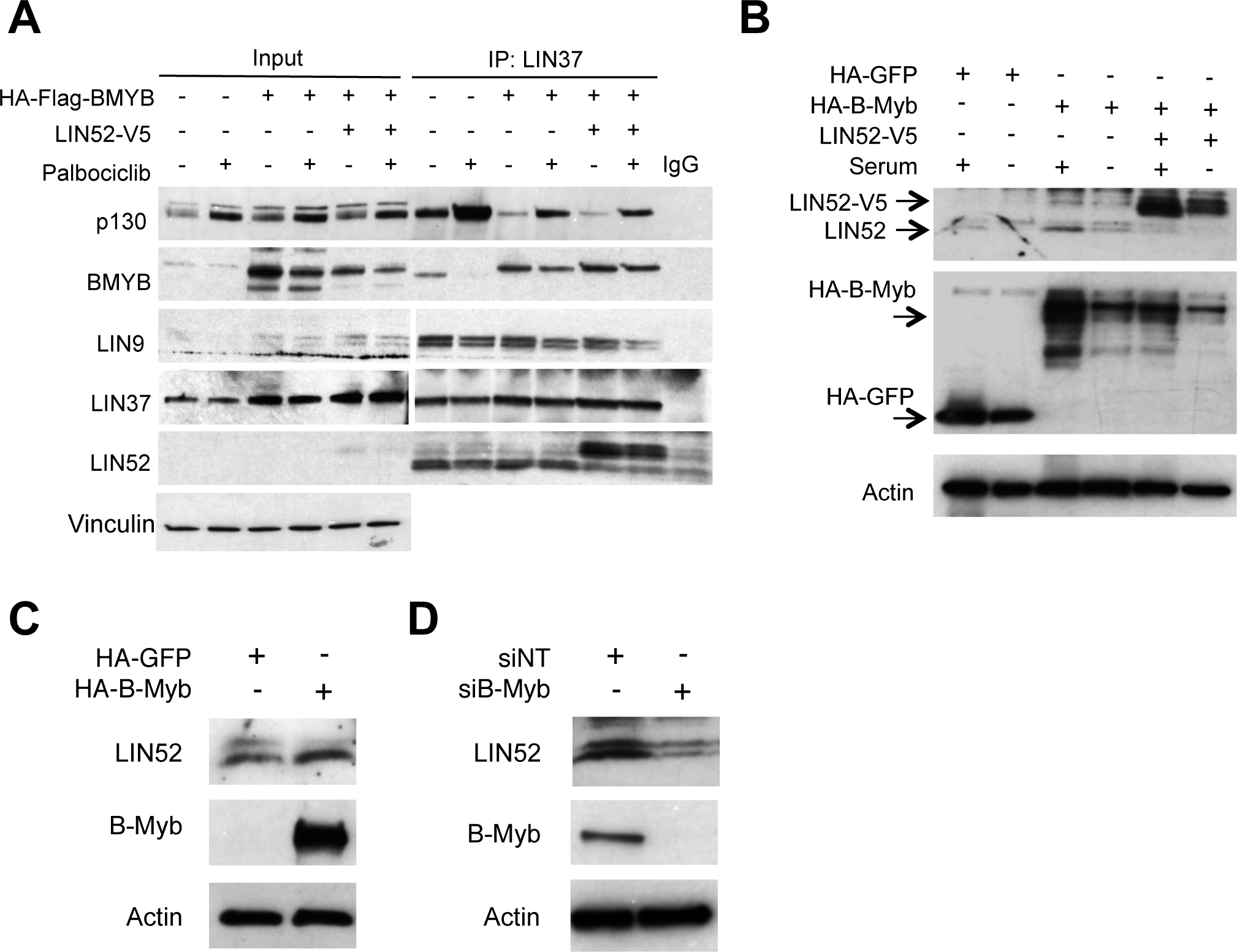
Co-expression of LIN52 fails to rescue DREAM formation in the presence of high B-Myb. (A) IP/WB analysis of the DREAM and MMB formation in BJ-hTERT stable cell lines expressing HA-B-Myb alone, or together with LIN52-V5, compared to parental cells. Cells were incubated with 10 μM palbociclib to induce G0/G1 arrest, or vehicle, for 18h before harvesting. Note that co-expression of LIN52-V5 does not rescue DREAM formation in B-Myb overexpressing cells compared to control. **(B)** Direct WB of the extracts from BJ-hTERT cell lines grown with or without serum for 24 h shows increased expression of endogenous LIN52 in BJ-hTERT cells expressing HA-B-Myb in the presence of serum, compared to control cells. Also, note decreased expression of endogenous LIN52 in the presence of ectopically expressed LIN52-V5. **(C)** WB analysis of the extracts from indicated BJ-hTERT cell lines shows lack of phosphorylated form of LIN52 in HA-B-Myb expressing cells. **(D)** WB analysis of the extracts from siRNA-transfected SKOV3 cells shows decreased expression of LIN52 in B-Myb-depleted cells compared to control.

To clarify this finding, we used the cell lines described in **Figure 2C** to detect changes in LIN52 protein by direct Western blot. Indeed, LIN52 was more abundant in HA-B-Myb-expressing cells in the presence of serum, and there was a decrease in the endogenous LIN52 protein levels when ectopic LIN52-V5 was expressed (**Fig. 3B**). Since *LIN52* is a DREAM target gene and high B-Myb expression is associated with DREAM disruption, it is possible that upregulation of LIN52 is due to transcriptional de-repression. In support of this model, we found that B-Myb and steady state LIN52 protein levels are tightly correlated. When B-Myb was overexpressed in BJ-hTERT cells, LIN52 was also more abundant, and the opposite effect was observed upon B-Myb knock-down in SKOV3 cells (**Fig. 3C, D**). Collectively, these results led us to investigate changes in LIN52 levels as a potential mechanism by which B-Myb impacts DREAM assembly.

We have previously characterized T98G cells stably expressing a LIN52 mutant S28A in which DREAM formation is diminished. ^14^ This substitution abolishes the DYRK1A-phosphorylation site essential for DREAM formation without interfering with the MMB complex assembly. ^15^ Interestingly, endogenous LIN52 protein was similarly down-regulated in T98G cell lines expressing the wild type LIN52-V5 or LIN52-S28A-V5, as compared to the empty vector control cells (**Fig. 4A**). Furthermore, RT-qPCR with primers designed for the 3’ untranslated (UTR) region shows no significant changes in endogenous LIN52 mRNA resulting from ectopic LIN52 expression, suggesting these observations are due to protein-level regulation (**Fig. 4B**). In support of this conclusion, ectopically expressed LIN52-V5 levels were tightly regulated by overexpression or knockdown of B-Myb in T98G cells (**Fig. 2S**).

**Figure 4:**
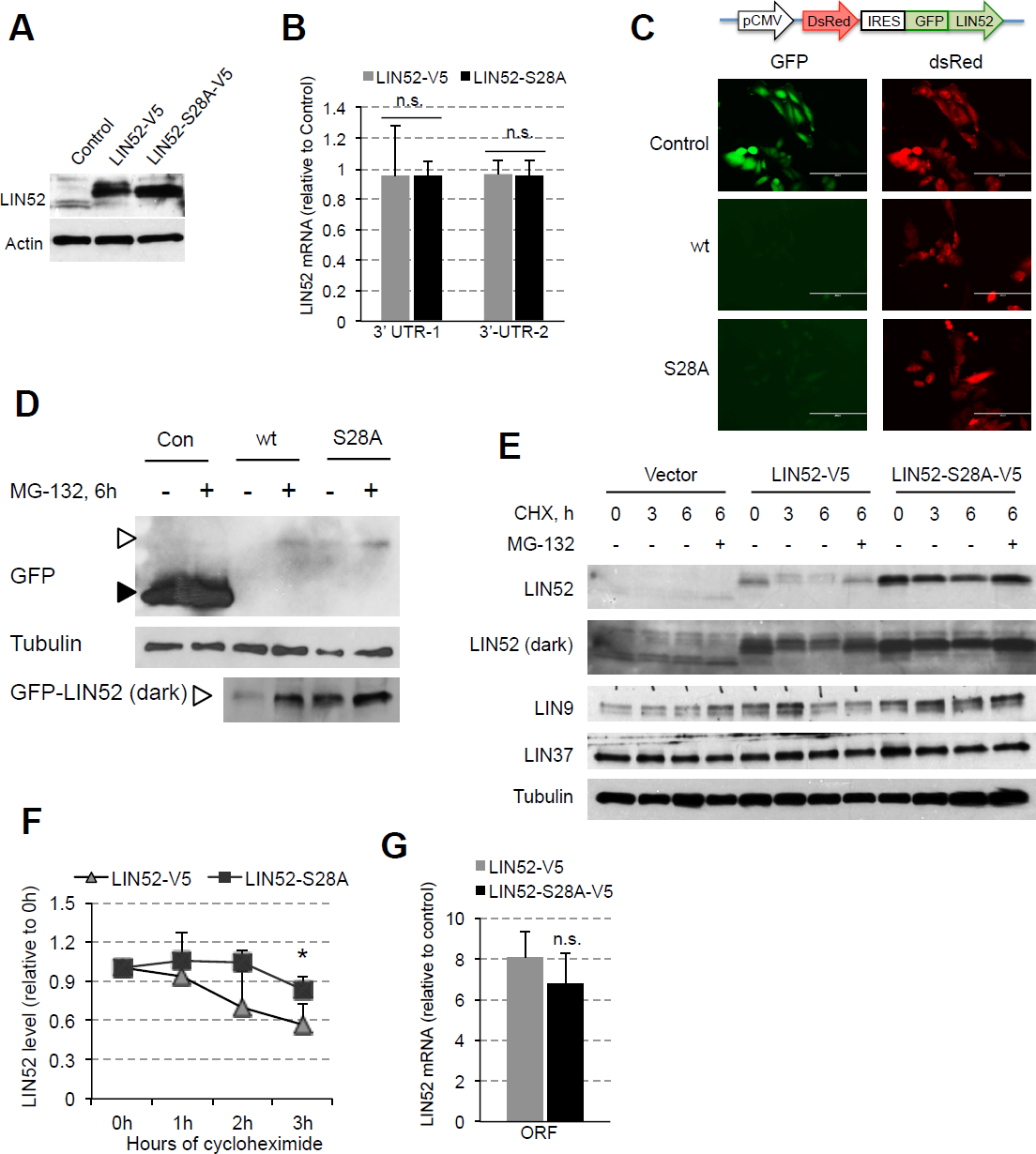
LIN52 is regulated at the protein level. (A) WB analysis shows decreased expression of endogenous LIN52 in T98G cell lines stably expressing either wild type, or S28A-LIN52. **B)** RT-qPCR analysis with primers designed to detect the 3’ untranslated region of endogenous LIN52 mRNA reveals no significant changes in the presence of ectopic LIN52. Graph shows average ± stdev (N=3, n.s. indicates p-value >0.05). Fluorescence cell imaging of T98G cells expressing GPS control, or GPS-LIN52 fusion constructs reveal decreased signal of GFP reporter fused to LIN52 compared to control, but not of the dsRed reporter protein. WB analysis confirms the expression of the GFP-LIN52 fusion proteins at a lower level than GFP alone. Cells were treated with MG-132 for 6h to inhibit proteasomal degradation. Note that S28A mutation increases the expression of the GFP-LIN52 fusion protein (but not to the GFP-control levels). **(E)** WB analysis of cycloheximide (CHX) chase experiment using T98G stable cell lines expressing empty vector, LIN52-V5 or LIN52-S28A-V5 mutant. One set of cells collected at 6h was treated with both CHX and MG-132 to assess proteasomal degradation. Note differential rates of degradation of LIN9, LIN37, as well as endogenous and ectopic LIN52. (**F**) Quantification of data from three independent CHX chase experiments using T98G cell lines expressing LIN52-V5 or LIN52-S28A-V5 mutant. Graph shows average ± stdev (* -Student’s t-test p<0.05). RT-qPCR analysis shows similar expression of the wild-type and S28A-mutant LIN52 mRNA. Graph shows average ± stdev of three independent experiments (ns -Student’s t-test p>0.05).

These findings prompted further investigations into LIN52 protein regulation using a Global Protein Stability (GPS) approach. ^36^ In this method, the protein of interest (LIN52) is fused with GFP and expressed as a single mRNA transcript together with an internal control marker (DsRed). During translation, an internal ribosome entry site in the GPS transcript allows for the production of LIN52-GFP and DsRed proteins (**Fig. 4C**). By monitoring the relative fluorescence intensities of GFP and DsRed, LIN52 abundance is measured in living cells. ^36, 37^ T98G glioblastoma cells were chosen for these studies as they have been used extensively in cell cycle research, including initial characterization of the DREAM complex. They also represent a cancer cell line lacking *MYBL2* amplification, making them amenable to both loss and gain of function studies of B-Myb. ^13, 14^, ^16, 38-42^ Using direct immunofluorescence and immunoblotting, we observed that although LIN52-S28A fusion protein was expressed at a slightly higher level than wild-type LIN52, both proteins were expressed at dramatically lower levels than control GFP alone. The dsRed expression levels were comparable in these cell lines, indicating that LIN52 fusion proteins are unstable. To determine whether LIN52 fusion proteins are degraded by proteasome, we treated the GPS-T98G cell lines with MG-132 for 6 hours. Addition of MG-132 increased LIN52-GFP abundance but had no effect on the corresponding S28A fusion, implicating LIN52 phosphorylation in the regulation of LIN52 protein level and the ubiquitin-proteasome pathway in LIN52 degradation (**Fig. 4D**). However, since inhibition of the proteasome by MG-132 or mutation of the serine 28 residue in LIN52 failed to fully restore LIN52 fusion protein level to that of the control GFP, it is likely that additional factors govern LIN52 level. Combined with our observations shown in **Figures 3C, D**, perhaps B-Myb is one of these factors.

We next directly compared the stability of the endogenous LIN52, wild-type LIN52-V5, and S28A mutant LIN52-V5 using cycloheximide (CHX) chase experiments with T98G stable cell lines. Quantification of three independent CHX chase experiments revealed a significant decrease in wild-type LIN52-V5 level relative to the S28A mutant after 3 hours of CHX treatment (**Figs. 4E, F**). Furthermore, although endogenous LIN52 was expressed at lower levels compared to the LIN52-V5 proteins, it showed greater stability than ectopically expressed wild type LIN52. As before, we noted a decrease in endogenous LIN52 protein levels resulting from its rapid degradation in the presence of ectopic LIN52 proteins. RT-qPCR analysis using primers designed to detect the LIN52 open reading frame (ORF) showed no significant difference between LIN52-V5 and LIN52-S28A-V5 mRNA expression (**Fig. 4G**), further confirming the predominance of protein level regulation. Additionally, we noted that while LIN52 was depleted over the course of 6h, two other MuvB proteins (LIN9 and LIN37) remained relatively stable (**Fig. 4E**). This result indicates that LIN52 could be differentially regulated among the other MuvB proteins.

Together, these results demonstrate that LIN52 is regulated at the protein level. Therefore, the changes observed in LIN52 level with depletion or overexpression of B-Myb (**Fig. 3B-D**) are likely due to protein-level regulation of LIN52 by B-Myb, and not by B-Myb-mediated disruption of the DREAM repression. Furthermore, these findings indicate that LIN52 level may be part of the mechanism by which B-Myb alters DREAM and MMB formation.

#### B-Myb regulates LIN52 level

To further investigate the role of B-Myb in regulation of LIN52, the effect of B-Myb expression on LIN52 protein stability was initially examined in a series of quantitative cycloheximide chase experiments. As shown in **Figures 5A and B**, LIN52-V5 protein levels were more rapidly diminished after 3 hours of CHX treatment in T98G cells after RNAi-mediated B-Myb knock-down compared to cells transfected with non-targeting (NT) control siRNA. Furthermore, over-expression of B-Myb in T98G cells resulted in significant stabilization of both ectopically expressed LIN52-V5 **(Figures 5C and D)** and endogenous LIN52 in BJ-hTERT fibroblasts (**Fig. 5E**). Interestingly, in the same series of experiments, we found that B-Myb exhibits no significant effect on LIN52-S28A-V5 protein level. Since B-Myb is still able to bind to LIN52-S28A-V5, these results suggest that phosphorylation at the serine 28 residue is also regulating LIN52 levels.

**Figure 5:**
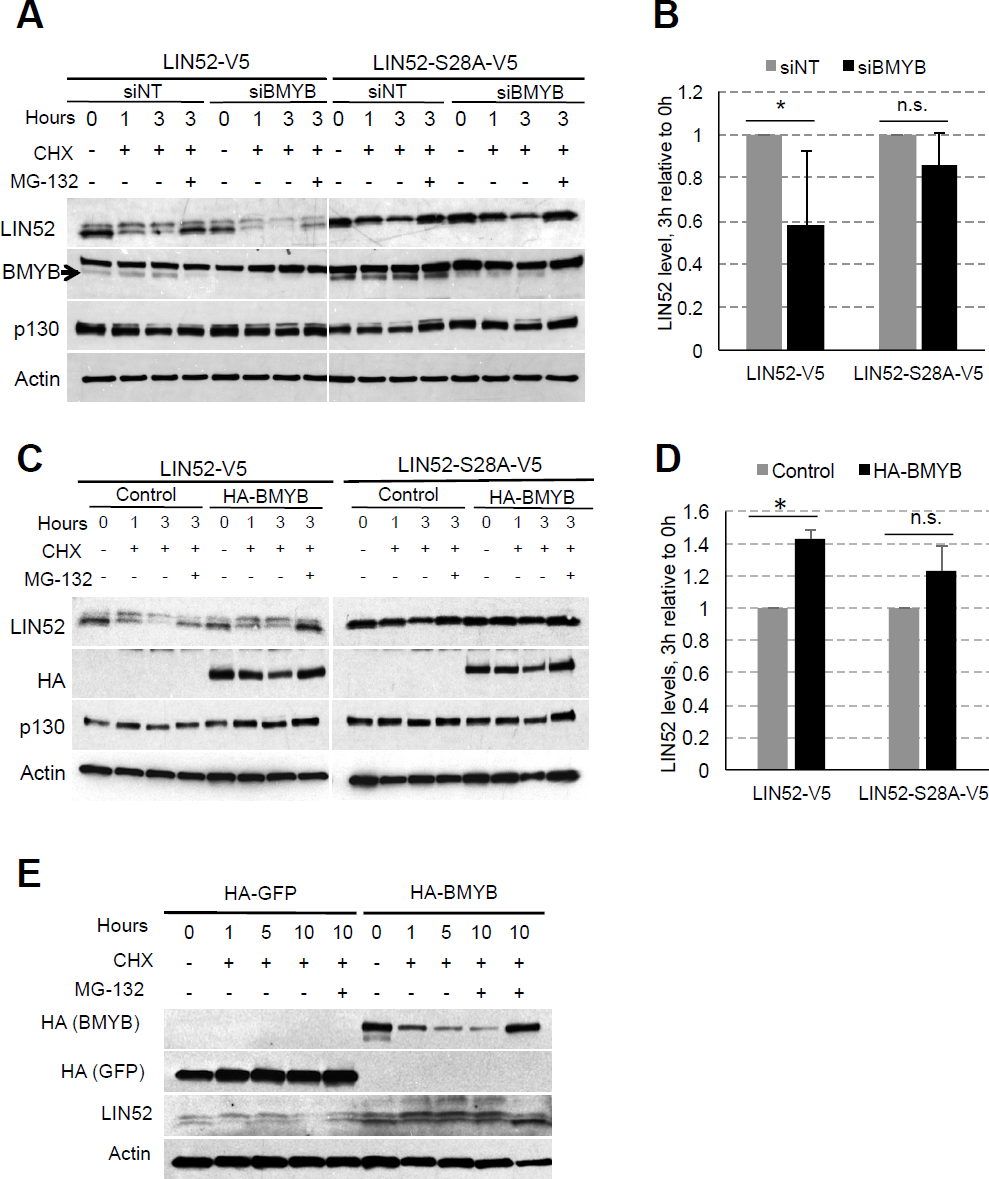
B-Myb regulates LIN52 protein turnover. (A) Representative immunoblot shows that RNAi knockdown of B-Myb decreases stability of the wild type, but not the S28A-LIN52. T98G cells stably expressing LIN52 proteins were transfected with siRNA and used for CHX chase assays after 36 hours. **(B)** ImageJ quantification of changes in LIN52 stability upon B-Myb knockdown. After normalizing to actin, the ratio of LIN52 band density after 3h of CHX treatment to 0h in si-B-Myb transfected cells was expressed relative to si-NT (control). Graph shows average ± stdev of three independent experiments shown in panel A (* and n.s. correspond to p-value <0.05 and >0.05 at 3h, respectively). Note that LIN52-S28A-V5 stability is not affected by B-Myb levels. **(C)** Representative immunoblot shows CHX chase assays using T98G cells stably expressing LIN52-V5 protein alone, or together with HA-B-Myb. **(D)** Quantification of the data shown in panel **C** was performed as described in **B. (E)** WB assay shows that stable overexpression of HA-B-Myb increases expression and stability of endogenous LIN52 in BJ-hTERT cells.

#### DYRK1A-mediated phosphorylation contributes to regulation of LIN52 stability

Since we previously demonstrated that LIN52 phosphorylation at serine 28 residue is mediated by DYRK1A, we used CRISPR/Cas9 to generate T98G cells devoid of DYRK1A protein (DYRK1A-KO) (**Fig. 4S**). Additionally, we treated intact T98G cells with a DYRK1A kinase inhibitor, harmine (10μM for 20h). As shown in **Figure 6A**, LIN52 was expressed at higher steady-state levels and appeared in a predominantly un-phosphorylated (faster migrating) form in DYRK1A knock-out cells or with harmine treatment, consistent with lack of DYRK1A activity (**Fig. 6A**). Furthermore, CHX chase experiments confirmed that stability of endogenous LIN52 was enhanced when DYRK1A is absent or inhibited by harmine (**Figs. 6B and C**). Interestingly, RT-qPCR analysis of the DYRK1A-KO or harmine-treated T98G cells revealed a modest increase in LIN52 mRNA expression when DYRK1A is knocked out or inhibited (**Fig. 6D**) suggesting that LIN52 is also regulated by DYRK1A at the level of transcription. To further determine the importance of DYRK1A in regulating LIN52 protein, we used RNAi to deplete DYRK1A in T98G cells expressing either V5-tagged wild-type or S28A mutant LIN52. We found that while a stabilization effect was observed in wild-type LIN52 upon DYRK1A knock-down, expression of S28A mutant was unchanged, further supporting the role of DYRK1A in regulation of LIN52 protein stability (**Figs. 6E and F**). Taken together, our data demonstrate that LIN52 is regulated at the protein level by B-Myb and DYRK1A-dependent phosphorylation, as well as at the transcriptional level by DYRK1A.

**Figure 6:**
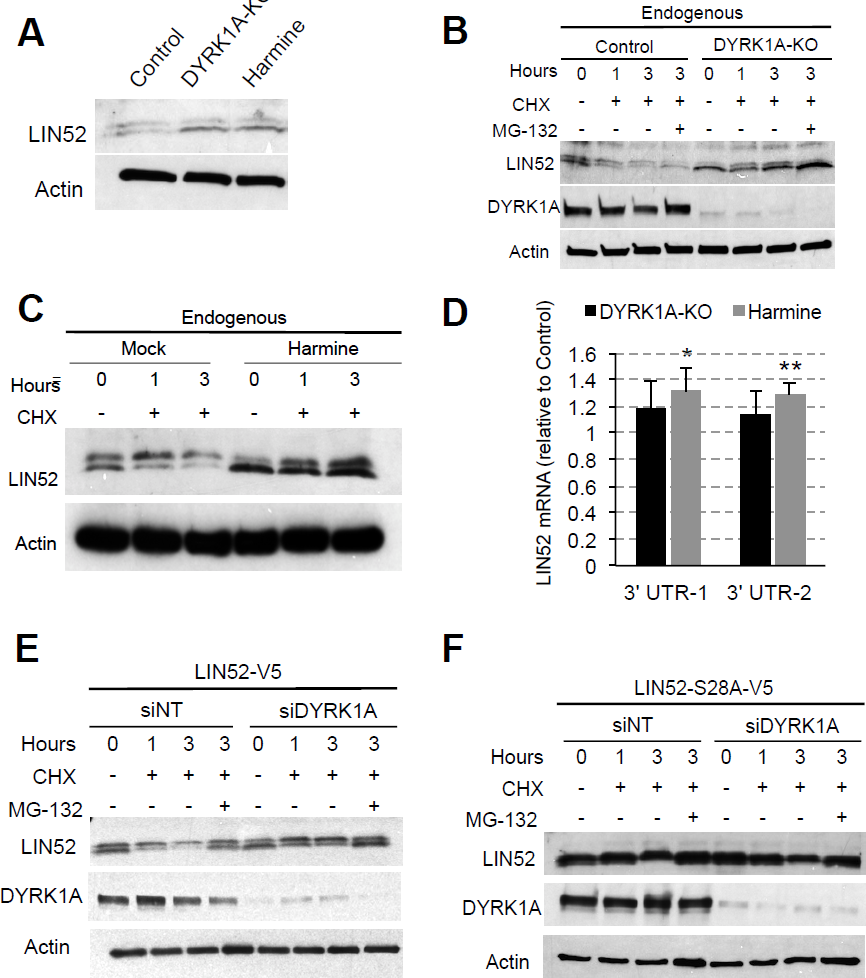
Inhibition of DYRK1A results in increased LIN52 expression. (A) Immunoblot shows increased steady-state LIN52 levels in T98G cells after CRISPR-Cas9 knock-out (KO) or inhibition of DYRK1A kinase, as compared to control cells. To inhibit DYRK1A, T98G cells were incubated with 10 μM harmine for 20h. **(B, C)** Endogenous LIN52 protein stability is increased upon harmine-induced inhibition of DYRK1A or in DYRK1A-KO cells, as compared to control. **(D)** RT-qPCR analysis of T98G cells shown in panel A reveals a modest but statistically significant increase in LIN52 mRNA expression when DYRK1A is inactivated, as compared to control cells. Graph shows average ± stdev of three independent experiments (* and ** correspond to p-value <0.05 and <0.01, respectively). **(E, F)** CHX chase assays reveal that transient si-RNA knockdown of DYRK1A results in increased stability of the ectopically expressed wild type, but not the S28A mutant LIN52.

#### B-Myb is highly expressed in cancers with LIN52 loss or DYRK1A gain

We hypothesized that high B-Myb expression is a compensatory mechanism for maintaining proliferation in tumors with LIN52 loss or DYRK1A gain. ^43, 44^ To test this model, high grade serous ovarian carcinoma TCGA data was queried for co-occurrence of *MYBL2* gene copy number gain with losses or gains in the *LIN52* or *DYRK1A* genes, and analyzed using cBioPortal.org tools. ^45^In support of our model, mutual exclusivity analysis showed a significant chance of co-occurrence of *MYBL2* copy number gain with either loss of *LIN52*, or gain of *DYRK1A*, in the same tumor sample (**Fig. 7A**). A similar analysis of the opposite scenario (*MYBL2* gain with *DYRK1A* loss or *LIN52* gain) found no significant associations. Overall, our data presented here supports a model in which the regulation of the MuvB core subunit, LIN52, could be important for proper switching between the DREAM and MMB complexes during the normal cell cycle. B-Myb amplification or over-expression in cancer could lead to stabilization of LIN52 and sequestering of the MuvB core away from the DREAM complex, resulting in preferential formation of the MMB complex under the conditions when LIN52 levels are low (**Fig. 7B**). This model could explain the observed increased expression of the DREAM and MMB target genes in breast and ovarian cancers with high levels of B-Myb. Furthermore, our model is mechanistically relevant to cancers in which DYRK1A kinase is active because B-Myb’s impact on LIN52 stability is less relevant when LIN52 cannot be phosphorylated by DYRK1A (**Figs. 5A-D**).

**Figure 7:**
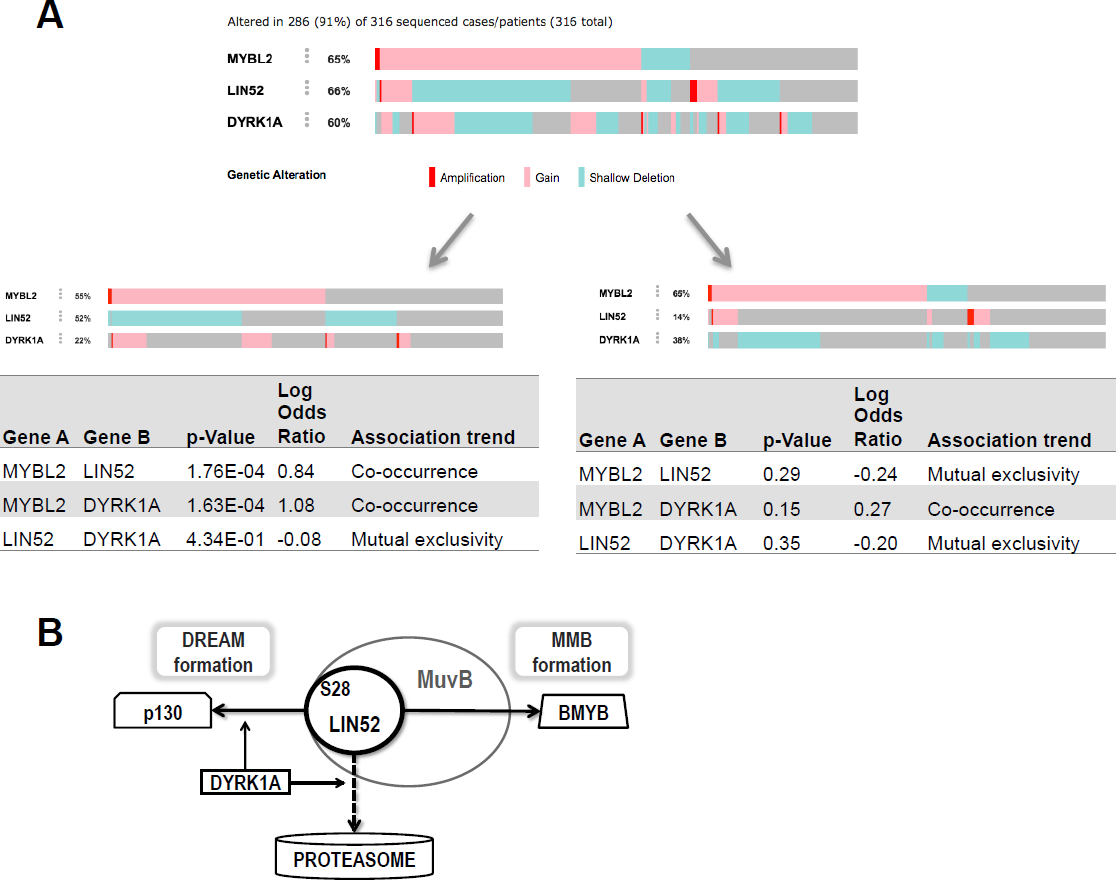
Model of DREAM disruption by high B-Myb. (A) Analysis of TCGA data using cBio.org Mutual Exclusivity tool shows that *MYBL2* amplification or gain is significantly associated with *DYRK1A* gain or *LIN52* loss in high-grade serous ovarian carcinoma. There is no significant association of *MYBL2* amplification or gain with *DYRK1A* loss or *LIN52* gain. **(B)** Schema of the proposed model, in which phosphorylation of LIN52 by DYRK1A kinase increases its turnover. When overexpressed, B-Myb stabilizes LIN52 and sequesters MuvB for increased MMB formation, resulting in cell cycle progression under the conditions favoring growth arrest.

## Discussion

Our data show that a gene expression signature of DREAM complex inactivation is present in breast and ovarian cancers with high expression of B-Myb, and that the DREAM complex is disrupted in human cells when B-Myb is highly expressed. B-Myb is overexpressed in many types of cancer and is recognized as a poor prognostic factor, but the known role of B-Myb as a transcription factor required for mitotic progression does not fully explain its established association with a more aggressive cancer phenotype. Furthermore, our studies presented here describe for the first time the complexity by which LIN52, the key MuvB component required for assembly of both DREAM and MMB, is regulated. LIN52 is tightly regulated at the protein level. Ectopic LIN52 expression results in increased turnover of both the endogenous and ectopically expressed protein. We also show that, while S28-phosphorylation-dependent proteasomal degradation plays a role in LIN52 protein turnover, this is clearly not the only mechanism that regulating its protein levels. Indeed, proteasomal inhibition or S28A mutation could only modestly increase the expression of the GFP-LIN52 fusion proteins in the GPS dual reporter system (**Fig. 4C**). Given a highly stable association of the MuvB subunits within the complex, it is possible that several MuvB subunits must be expressed together so they can be properly folded and stabilized. It will be interesting to determine whether this additional, phosphorylation-independent mechanism could involve cellular chaperone complexes, such as CCR4-NOT, that mediate degradation of misfolded proteins or products of aberrantly-expressed mRNA. Perhaps these cellular chaperone complexes play a role in maintaining the proper levels of MuvB core subunits in the cell.

Several earlier studies have shown that DREAM and MMB complexes exist during different stages of the cell cycle with minimal overlap during the G1-S transition. However, until now, it was not clear whether the MMB formation simply occurs when the MuvB core is released from DREAM due to increasing cyclin-CDKs phosphorylation of the DREAM components, or whether B-Myb plays a active role in DREAM disassembly. Our data allows consideration to a model in which the initial expression of B-Myb could occur as a result of DREAM phosphorylation by CDKs, followed by B-Myb actively sequestering the MuvB core away from DREAM. Further studies using B-Myb mutants deficient in binding to MuvB, or MuvB subunit mutants that can not interact with B-Myb, will help to test this model. Furthermore, we characterize the mechanism by which stabilization of LIN52 at the protein level by B-Myb could contribute to the sequestering MuvB away from DREAM formation. This mechanism could be particularly important under conditions when LIN52 level is kept in check by DYRK1A kinase activity. Interestingly, we noted that overexpression of B-Myb in non-transformed BJ-hTERT cells reduced the appearance of the phosphorylated form of LIN52, consistent with the increased stability of un-phosphorylated LIN52 protein (**Figs. 3B and C**). These findings could suggest another potential mechanism in which B-Myb stabilizes LIN52 by preventing its phosphorylation at S28. In support of this model, we have shown that, contrary to our observations with wild-type LIN52, changes in B-Myb level do not impact on the stability of LIN52-S28A (**Figs. 5**). Therefore, B-Myb can stabilize LIN52 by protecting it from proteasomal degradation though steric hindrance, or by blocking access of DYRK1A to the S28 site in LIN52. Additionally, B-Myb could stabilize LIN52 by recruiting a phosphatase that will remove the LIN52 phospho-S28 degradation mark. In addition to physical interaction between LIN52 and B-Myb, transcriptional activity of B-Myb is also an important consideration. For example, B-Myb might regulate expression of proteins that could stabilize LIN52. Therefore, further studies are needed to investigate these possible mechanisms.

We also demonstrate that B-Myb overexpression leads to de-repression of DREAM cell cycle gene targets (**Figs. 1D, 2B**). Since *MYBL2* itself is a DREAM target gene, this might create a positive feedback loop in which B-Myb overexpression drives further B-Myb expression and perpetuates DREAM disruption. ^46^ This is also relevant to HPV-positive cancers with E7-mediated DREAM complex disassembly. ^46-50^ Indeed, T98G cells expressing viral oncoproteins have higher amounts of B-Myb and MMB as compared to cells with intact DREAM. ^51, 52^ B-Myb was also identified as a target for E7-mediated transcriptional activation, leading to inappropriate expression of B-Myb in G1 and higher steady state levels in cycling cells, mimicking our model of B-Myb overexpression. ^53^ Clinical evidence supports these studies by showing that E7 expression correlates with B-Myb levels and mitotic gene expression in cervical cancer. ^54, 55^ Together, these studies provide a support to mechanisms whereby E7 and B-Myb cooperate to increase MMB formation and mitotic gene expression at the expense of pocket protein/DREAM activity. Furthermore, E7 has been shown to disrupt the interaction between LIN52 and the p130/p107, supporting the importance of LIN52 in mediating DREAM and MMB formation. ^56^

LIN52 is required for both DREAM and MMB formation and directly interacts with p130/p107 *in vitro,* but is insufficient for co-precipitation of B-Myb. Therefore, other MuvB components are also important for mediating MMB formation and perhaps LIN52 mainly serves to preserve MuvB integrity. ^15^ This is supported by diminished co-immunoprecipitation of MuvB complex components when LIN52 is depleted by shRNA. ^14^ However, other studies have shown that B-Myb, LIN37, and LIN54 are also diminished when LIN9 is depleted, suggesting that there is no significant free pool of these proteins and they form a stable complex in the S phase. ^57^ Another recent study revealed compromised DREAM repressor function upon LIN37 loss, but MMB function remained intact and the remaining MuvB subunits were able to interact with either p130, or B-Myb in the LIN37 knockout cells. A MuvB-binding deficient mutant LIN37 was unable to rescue DREAM, demonstrating the importance of MuvB integrity for DREAM assembly. ^58^ In combination, these findings highlight the complexity of MuvB regulation and suggest that MuvB components may be differentially regulated depending on their participation in either DREAM or MMB. We have shown that LIN52 levels in the cell are tightly controlled and regulated by B-Myb and DYRK1A. Together, these findings suggest that regulation of LIN52 stability may be central to maintaining MuvB and alterations in LIN52 stability, in turn, a potential determining factor for which complex (DREAM or MMB) forms. We have previously shown that B-Myb and p130 can bind MuvB simultaneously, so direct competition for binding is unlikely. ^15^ However, if LIN52 is crucial to the integrity of MuvB, then B-Myb may hold a competitive advantage over p130 through its stabilization of LIN52 (**Fig. 5**).

The same phosphorylation event resulting in DREAM assembly is also associated with LIN52 degradation (**Fig. 6**). ^14^ This suggests LIN52 degradation is a cooperative process between the DREAM complex and another factor, such as an E3 ligase. Previous proteomics data identified several candidate E3 ligases associated with the DREAM complex that may play a part in LIN52 degradation. ^13, 14^ Determination of the particular E3 ligase responsible for LIN52 degradation would be valuable to further understand the regulation of the DREAM and MMB complexes. ^8^ Finally, several of the MMB downstream targets upregulated in late S-G2 phases of the cell cycle, such as Aurora kinase A, Polo-like kinase B and others have been proposed for developing anti-cancer therapies. ^8, 59^ Better understanding of the mechanisms that bring about high expression of these genes in cancer will be important to inform the future pre-clinical and clinical studies and to optimize patient stratification. Overall, we have shown a novel model describing some of the molecular processes underlying deregulated cell cycle gene expression in cancers with B-Myb amplification or overexpression. These findings argue that B-Myb is not only a negative prognostic factor through increased MMB formation and activation of the mitotic gene expression program, but also through decreased DREAM assembly and loss of repression of more than 800 cell cycle-regulated genes. Further studies with tumor samples are needed to validate our model in patients.

## Acknowledgements

We thank Ross Mikkelsen and Steven Grossman for critical reading of the manuscript and Sophia Gruszecki for her technical support. This project supported in part by R01CA188571 (L.L.).

## Methods

### Cell lines

Parental cell lines were purchased through the American Type Culture Collection. T98G cells stably expressing V5-tagged LIN52 or LIN52-S28A were previously produced by retroviral gene transfer. ^13, 14^ These cells were, in turn, infected with a HA-FLAG-B-Myb pMSCV-NTAP retroviral vector made previously by Gateway cloning and selected in media containing 1ug puromycin. BJ-hTERT cells were infected with the same HA-FLAG-B-Myb pMSCV-NTAP retroviral vector or a matching GFP-HA-FLAG control vector. ^13^ BJ-hTERT cells co-expressing both HA-FLAG-BMYB and LIN52-V5 were made in the same manner as the corresponding T98G cells described above. The LIN52-GFP and LIN52-S28A-GFP constructs were generated in GPS-pHAGE vectors and subsequently used to produce stable T98G cell lines.

T98G DYRK1A KO cells were established using GeneArt CRISPR Nuclease vector with OFP reporter (Life Technologies) harboring a DYRK1A-specific guiding RNA sequence (top strand: 5´-TCAGCAACCTCTAACTAACC-3´, bottom strand: 5´-GGTTAGTTAGAGGTTGCTGA-3´). T98G parental cells were transfected with control or DYRK1A gRNA-CRISPR plasmids, FACS-sorted for OFP expression, and plated onto 96-well plates to obtain single-cell clones. Individual clones were then screened for DYRK1A expression using immunoblotting. DYRK1A expression was also analyzed in subsequent early passages. Genomic DNA from clones with no detectable DYRK1A expression was isolated and used for PCR amplification of the DYRK1A genomic region 400 bp up-and downstream of the Cas9 targeting site using nested primers (first set: forward 5´-AAACCTGGCAGCAGGTGC-3´ and reverse 5´-CTCATCACACATCAAATATCCG-3´; second set: forward 5´-TGAGTTGGTTTGTGAGGAGC-3´ and reverse 5´-ACTTTCACACAACTACAGC-3´). PCR products were sequenced to confirm the presence of mutation at the targeted site. Since we received sequences with multiple lesions, we wanted to rule out the presence of a mixed clone and to confirm the presence of single cell clones. TA cloning was performed using Promega pGEM^®^-T Easy Vector Systems followed by transformation according to manufacturer’s protocol. White colonies (n=21) were isolated for each clone, allowed to grow on 96-well plates overnight, replica-plated and sent for hi-throughput sequencing using a T7 primer.

### Antibodies

Antibodies for MuvB complex components LIN9, LIN37, and LIN52 (Bethyl Inc.) were previously described in Litovchick et al. 2007. ^13^ Antibodies against tubulin (Sigma, SAB1411818), β-actin (Cell Signaling, 4970S), vinculin (Sigma, V9131), GFP (Santa Cruz, sc-9996), p130 (BD Biosciences, 610262), B-Myb (Santa Cruz, SC-724 [N-19] and Millipore, MABE886, clone LX015.1), DYRK1A (Cell Signaling, 3965), V5 (Bio-Rad, clone SV5-Pk1), and HA (Cell Signaling, 3724S) are commercially available. For immunoprecipitations using LIN37 (Bethyl Inc.), Reliablot^®^ Western blot kit (Bethyl Inc.) was used for subsequent probing with rabbit antibodies.

### RNA interference and transient transfections with plasmid DNA

Invitrogen RNAiMAX^™^ was used for all transfections of siRNA oligonucleotides according to the manufacturer’s instructions. Ambion^®^ Silencer^®^ Select MYBL2 human (s9119) and DYRK1A human (s4399) siRNA oligonucleotides were used. Transient transfections of plasmid DNA were performed using polyethylenimine, as described in Longo et al. 2013. ^60^

### Growth assay

ATPLite 1,000 assay kit (PerkinElmer, 6016941) was used according to the manufacturer’s protocol. For each cell line, 3,000 cells were plated per well and luminescence was measured at days 3 and 5.

### Real-time and quantitative PCR

RNA extractions were performed with TRIzol reagent (Invitrogen) and was reverse transcribed using SensiFAST^™^ cDNA Synthesis kit (Bioline). The resulting cDNA was used for qPCR with Maxima SYBR Green/ROX Mastermix (2X) (ThermoFisher, K022) and the following primers:

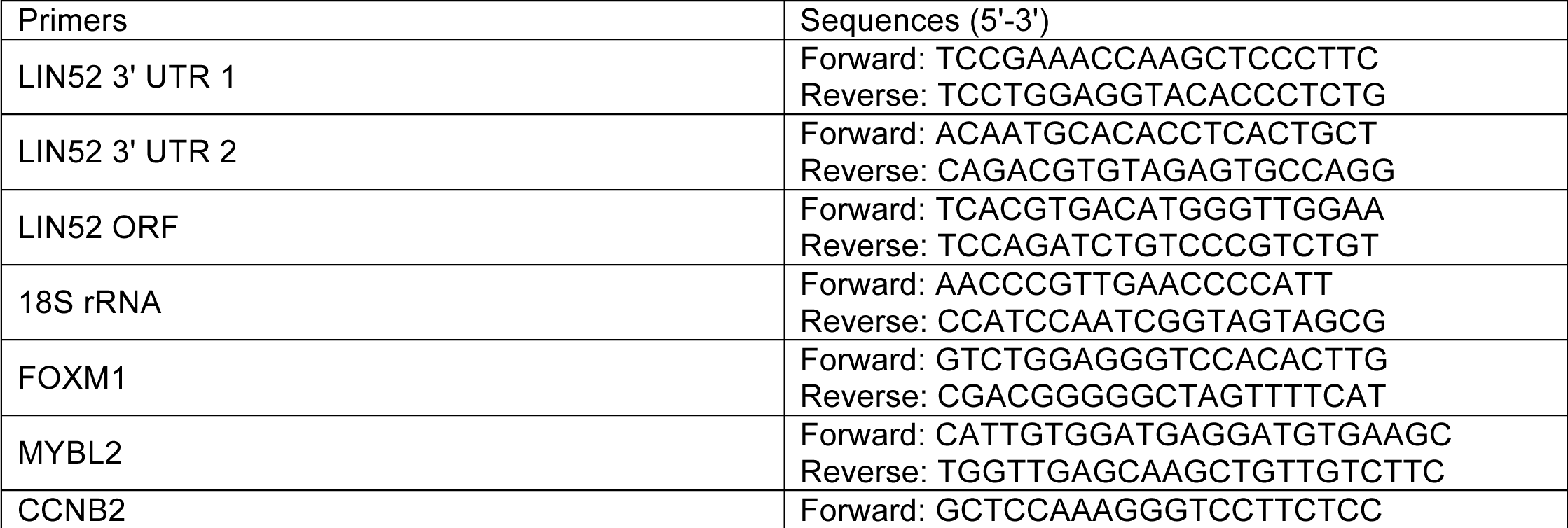
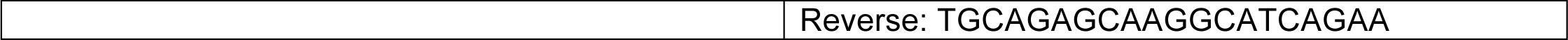

Applied Biosystems 7900HT was used for detection and fold changes in mRNA expression relative to controls were calculated using the 2DDCt methodology.

### Drugs

The following were obtained from Sigma-Aldrich: Cycloheximide (3-[2-(3,5-Dimethyl-2-oxocyclohexyl)-2-hydroxyethyl]glutarimide, Actidione, Naramycin A, C7698), MG-132 (Z-Leu-Leu-Leu-al, C2211), Harmine (7-Methoxy-1-methyl-9H-pyrido[3,4-b]indole, 286044), Palbociclib (PD 0332991 isethionate, PZ0199).

### Biostatistics

TCGA data analysis: Open access gene expression data summarized as RSEM values was obtained using the TCGA2STAT R package v. 1.2, along with the corresponding clinical annotations. Data for each cancer were obtained separately. The data was log2-transformed.

Differential expression analysis: Samples in the selected cancer cohort were sorted by expression of the selected genes. Differentially expressed genes were detected between samples in the upper 75 percentile of the expression gradient and samples in the lower 25 percentile using the limma R package v. 3.32.6. P-values were corrected for multiple testing using False Discovery Rate (FDR) method. Genes differentially expressed at FDR < 0.01 were selected for further analysis.

Functional enrichment analysis: Genes up-and downregulated in the selected cancer were overlapped with DREAM and MMB targets. ^13, 54^ Functional enrichment analysis of these gene lists was performed using Metascape (http://metascape.org/, [PMID: 26651948]) using the latest 03-16-2017 database version.

**S.Figure 1.**
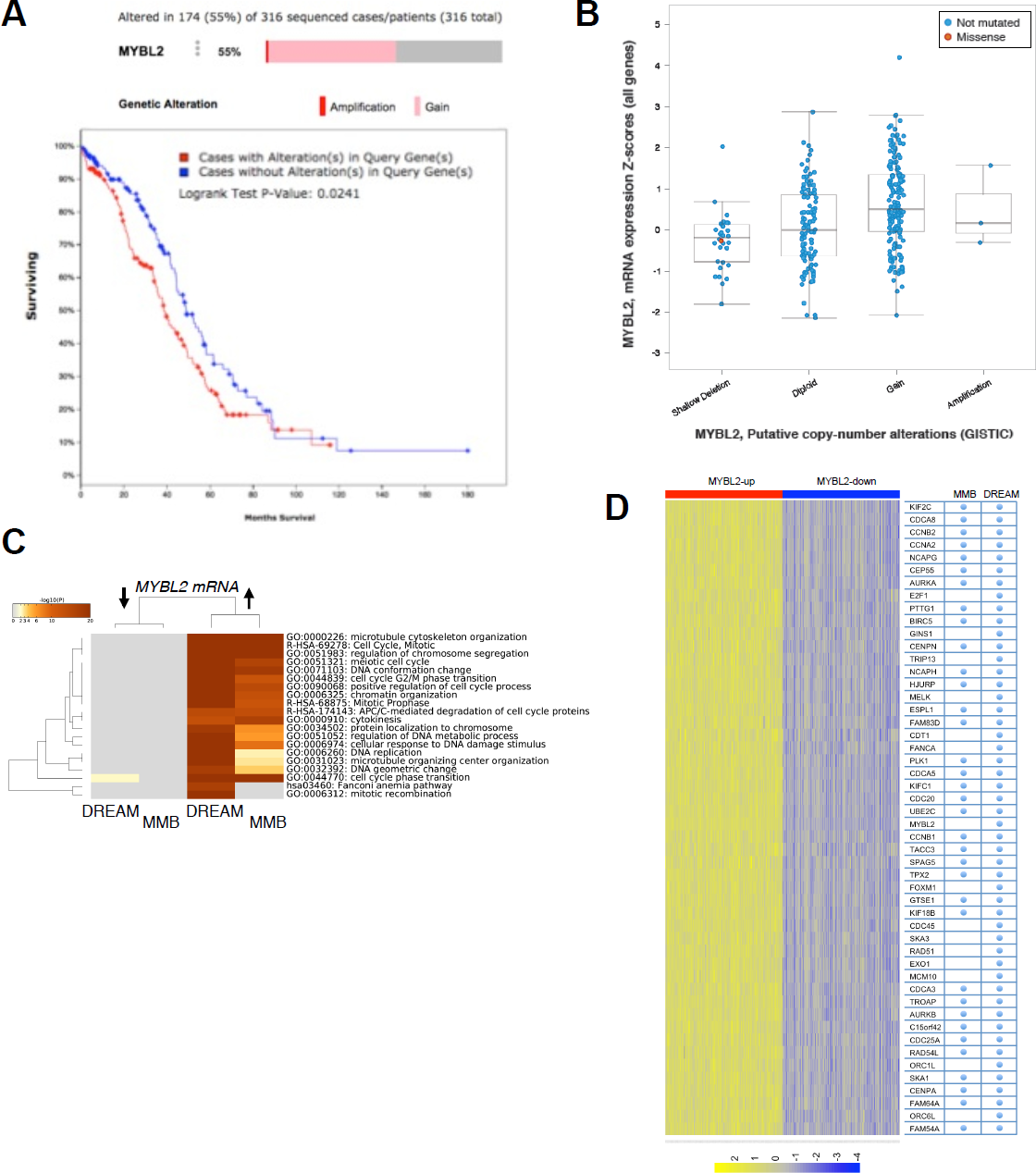
Supplemental Figure 1. (A, B) Analysis of TCGA data using cBio.org tools shows that gain of the *MYBL2* gene is associated with decreased survival in high-grade serous ovarian cancer patients, and that gain of *MYBL2* gene in HGSOC is associated with increased expression of its mRNA. **(C)** DREAM and MMB target genes are significantly upregulated in breast cancers with high B-Myb expression (Fisher’s exact test p-value 1.242E-102). **(D)** All 50 top up-regulated genes in our TCGA gene expression analysis of breast cancer tumors with high expression of B-Myb were identified as DREAM targets (χ^2^ with Yates correction p<0.001).

**S.Figure 2:**
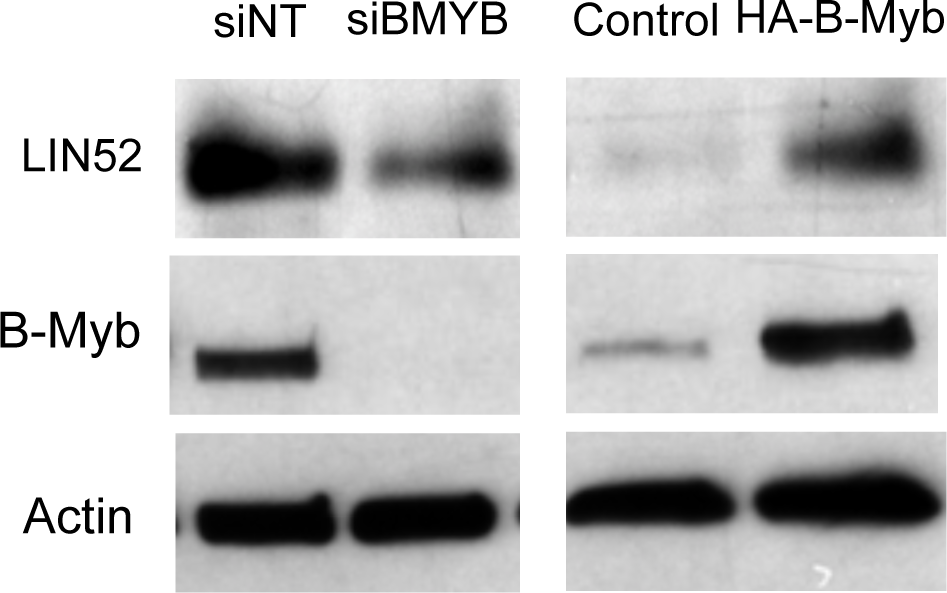
Supplemental Figure 2: (A) Steady state levels of stably expressed LIN52-V5 protein correlate with B-Myb expression in T98G cells. **(B, C)** RT-qPCR analysis reveals no significant changes in the endogenous (3’ UTR) and ectopic LIN52 mRNA (ORF) levels upon B-Myb knockdown (**B**), or overexpression (**C**) in T98G stable cell lines expressing LIN52-V5 proteins. Graphs show average ± stdev (N=3, n.s., p-value >0.05).

**S.Figure 3:**
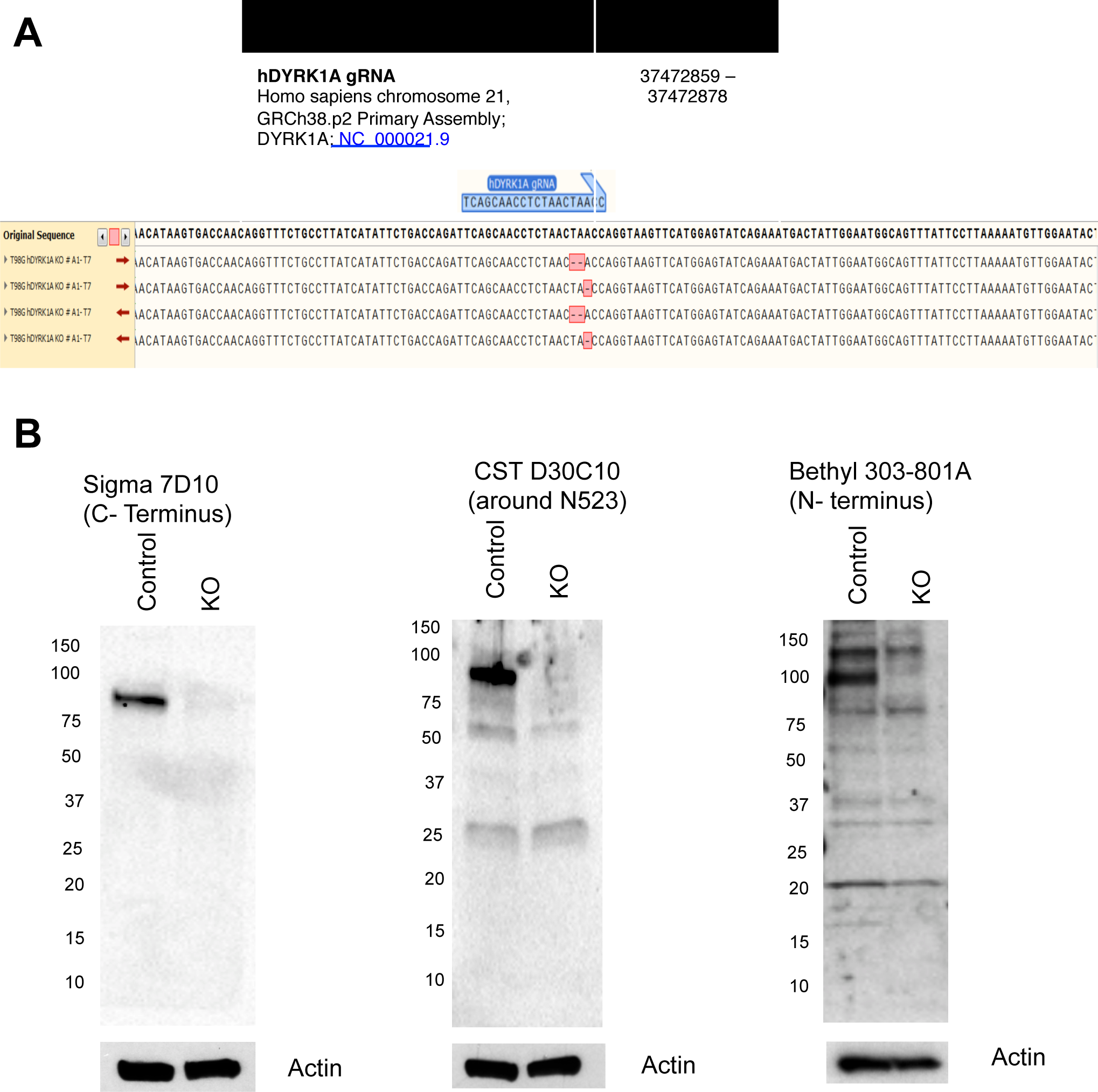
Supplemental Figure 3: (A) Genomic DNA position of DYRK1A and the gRNA binding coordinates used for generation of DYRK1A null cells by CRISPR/Cas9. **(B)** Sequences obtained from four different clones after TA cloning. Note that approximately 50% of the sequences of the 21 clones sequenced harbored the same mutation suggesting that each allele carried a different mutation. **(C)** Lysates from control (T98G parental cell line) and T98G KO cells were analyzed by immunoblotting using the indicated DYRK1A antibodies. Actin was used as a loading control.

